# Stem water content is crucial to support fruit tree functioning during heatwaves in a Mediterranean climate

**DOI:** 10.1101/2024.09.29.615635

**Authors:** Laura Rez, Justine E. Missik, Gil Bohrer, Yair Mau

## Abstract

Droughts are expected to intensify in the Mediterranean region due to climate change, yet the effect of these highly variable events on local trees is unknown. To study the particular effect of heatwaves in orchards, where soil-drought can be mitigated by irrigation, we propose a heatwave definition that focuses on atmospheric stress and its consequences, by relating the intensity of high VPD events to losses in tree stem-water storage (StWS). We found that the sensitivity and resilience of StWS to heatwaves is species-specific, and varies among species with different water-management strategies (e.g., isohydric orange and anisohydric mango trees, *p* < 10^−3^). Navel orange trees were sensitive to heatwaves starting at the 80th percentile of VPD in early spring, and once irrigation began, despite the harsh Mediterranean summer temperatures, StWC increased to 0.57 g cm^−3^, slightly greater than the StWC of the earlier wet season (approximately 0.55 g cm^−3^). Oppositely, there was a net reduction in StWC in Shelly mango trees from 0.75 to 0.69 g cm^−3^ between the two seasons, as sensitivity to heatwaves increased from the 90th to the 80th percentile in spring and summer, respectively. By first quantifying heatwaves and relating this new variable to changes in StWS, we were able to describe the sensitivity of each species according to the rarity of the heatwave events by VPD percentile, and their resilience to heatwaves over seasons based on the corresponding net changes in StWC. Though the experiment in this study was performed in a Mediterranean climate, hotter-droughts are rising globally and the framework developed here for quantifying and measuring the effect of heatwaves can be broadly applied across geographic locations.

## 1 Introduction

Climate change is driving a global intensification of the water cycle (Christian et al., 2021; Huntington, 2006), which is responsible for an increase in drought occurrence and intensity (Yuan et al., 2019; Rauscher et al., 2008; Sung et al., 2023). Drought and heatwave intensification has been observed in many regions from Asia to the Americas. The response of trees to the increasing recurrence, intensity and duration of dry heatwaves, whether in natural or man-made landscapes across the globe, is still largely unknown. In the Mediterranean region, the harsh combination of a growing dry season, intensifying heatwaves, and pest outbreak has already begun to shift the landscape from forest to shrub- and grass-land (Lloret and Batllori, 2021), and forest area is expected to continue to decline according to climate projections for the century (Hammond et al., 2022). Even in Mediterranean orchards where trees are irrigated and sprayed, heatwaves may pose a serious threat to tree water status, productivity, and product quality, of both historically-established species and recently adapted tropical fruit trees (Quiller et al., 2017; De Ollas et al., 2019; Fraga et al., 2020; Lorite et al., 2020; Scuderi et al., 2022). While numerous, local fruit-tree cultivars have demonstrated drought tolerance in studies of deficit irrigation, where similar or improved yield was found at the cost of (unnecessary) vegetative growth (García-Tejero et al., 2011; Panigrahi et al., 2014; Pleguezuelo et al., 2018), their fate facing intensifying atmospheric droughts in the region is unknown, and is particularly concerning with temperatures consistently soaring above 40 ^*°*^C in recent years.

To study tree sensitivity to dry heatwaves, we adopt the approach defined by Smith (2011) that an extreme weather event (EWE) should be tied to the statistical rarity in both the driver and the response; in the context of our study, this would refer to a distinct physiological response in the tree. Therefore, an appropriate definition of a heatwave will be one which is formulated regarding the statistical rarity of the relevant climate driver(s) (such as temperature and/or VPD) and physiological response(s) from the tree that are dependent on these drivers (such as changes in the water status or productivity of the tree). This is a simple but crucial distinction that will help to develop a robust definition of a heatwave in the context of tree studies, as until recently, EWE definitions have often varied by study and have received criticism for being arbitrary (De Boeck et al., 2010), inconsistent, and lacking statistical significance (Slette et al., 2019).

While such a specific definition of a heatwave may not yet exist, the general phenomena and related physiological responses, themselves, have been well-documented. Heatwaves have been linked to changes in a tree’s productivity, conductivity, water storage, and overall water-use efficiency (Teskey et al., 2015; Tatarinov et al., 2016; Urban et al., 2017; Birami et al., 2018; Drake et al., 2018; Salomón et al., 2022; Neuwirth et al., 2021). In central Israel, which experiences a Mediterranean climate, dry heatwaves are commonly experienced throughout the year, and especially in the six-month dry-season spanning spring to autumn (Shachak et al., 1998; Qin et al., 2006). Recently, it has been shown that trees in this environment prevent significant and potentially mortal drops in leaf-water potential by buffering high early-morning to afternoon transpiration demand with internal water storage (WS) from both the foliage and the stem (Preisler et al., 2022), illustrating the quick response from internal water storage to changing climatic conditions that drive transpiration (namely, vapor-pressure deficit (VPD).

Stem water storage (StWS) likely possesses a central role in this buffering, given that the stem contains the majority of tree WS (Köcher et al., 2013; Waring and Running, 1978) and may therefore further reflect the additional impact of a heatwave (when characterized by high VPD (Merilo et al., 2018)) on transpiration. This potential relationship, however, is not quantifiable given current heatwave definitions. Existing definitions, though varying in exact method, are particularly limited to identifying heatwaves according to a climate driver surpassing a defined threshold (Schär et al., 2004; Urban et al., 2017; Drake et al., 2018; Sun et al., 2014; Cochavi et al., 2021). Figure 1 exemplifies three of these fixed absolute-threshold definitions. An advantage of quantifying heatwaves using magnitudes on a continuous scale, as opposed to a binary classification, is that a continuous classification allows us to quantify the relationship between changes in StWS as a response to heatwaves using a continuous response curve, and to more broadly describe a tree’s sensitivity or resilience to these extreme weather events.

**Figure 1:**
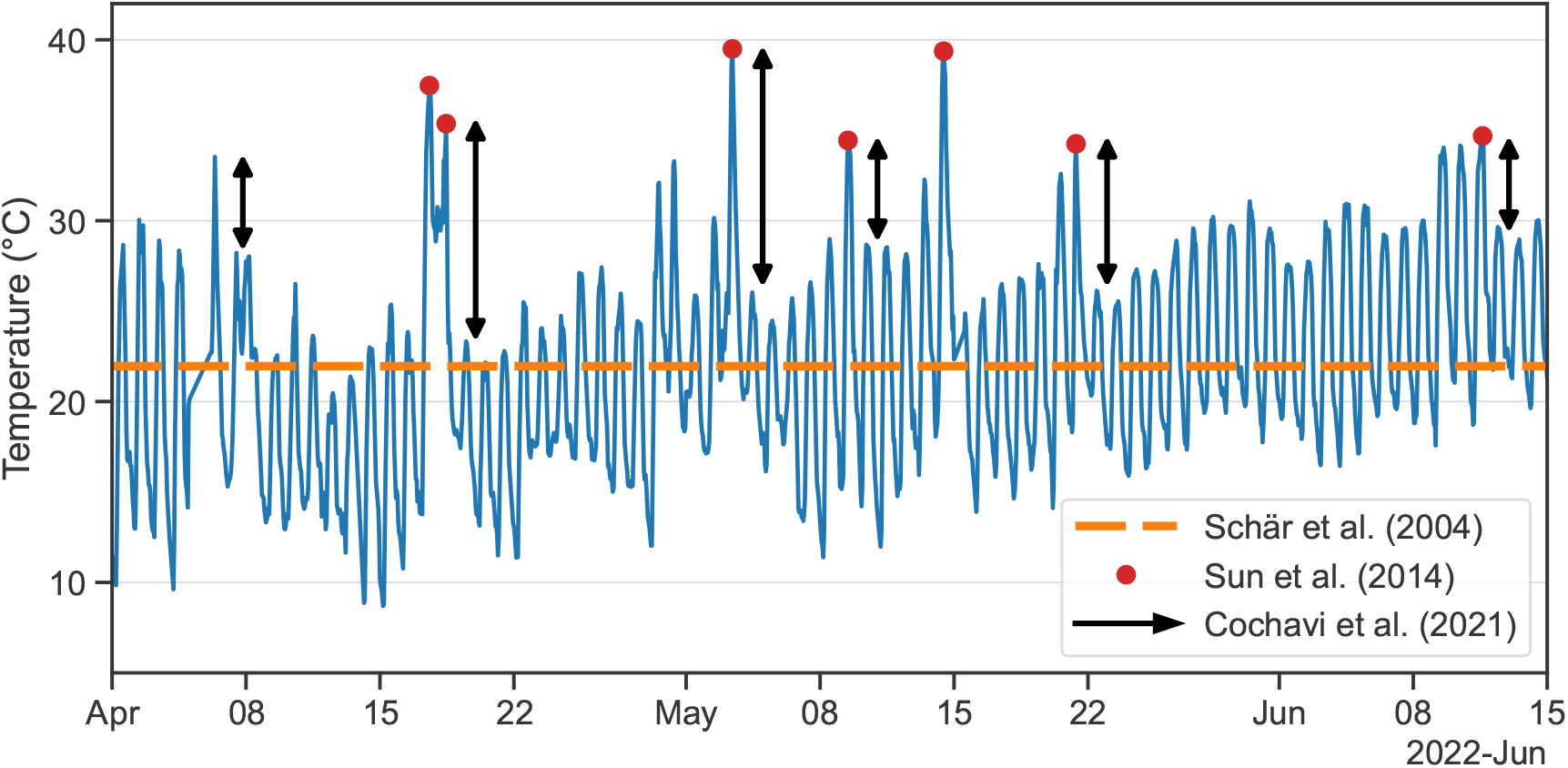
Various interpretations of climate dynamics for defining a heatwave, demonstrated on air temperature measured at the research site over the study period. The dashed orange line is the mean temperature over the study period, which, if greater than a historical mean by a specified number of degrees Celsius, constitutes a heatwave (Schär et al., 2004); the red circles indicate the maximum daily temperature that surpasses a defined threshold (corresponding to events as defined by Sun et al. (2014), for a threshold of 35 °C); the black arrows indicate the value of the difference between two days of the specified climate variable exceed a defined thresholds (corresponding to heatwaves as defined by Cochavi et al. (2021), for a temperature differential threshold of 5 °C)

In this study, we first present a theoretical relationship between heatwaves (statistically high VPD events) and changes in StWS related to these events, derived from the relationship between transpiration and VPD. We define the magnitude of a dry heatwave as the product of its intensity (VPD above a significant percentile threshold) and the duration of time it remains at this intensity. As this quantification yields a new unit for the magnitude of a heatwave (the product of VPD and time), we have given it a name for reference, Cumulative Heatwave Unit (CHU, measured in kilopascal hours). We hypothesize that heatwave magnitude is proportional to the loss in StWS above a typical diurnal baseline, caused by the heatwave. Our goal was to compare this theoretical definition of a dry heatwave, which is rooted in climate driver and tree response variability, with observed climate and StWS measurements from two tree species with opposing water management strategies and history in the region: relatively isohydric orange trees, which are a long-standing crop adapted to the region, and relatively anisohydric mango trees, which are a more recently introduced crop that are highly suited to the local winter-spring environment.

## 2 Materials and Methods

### 2.1 Research site

The research site is located in the Joseph Marguleas Experimental and Demonstration Farm of the Hebrew University, Faculty of Agriculture, Food and Environment, in Rehovot, Israel (34°47’43.3”, 31°54’14.9”). The study period is from March 22nd to June 30th, 2022, spanning early rain-fed spring to early dry summer, with irrigation beginning on April 26th. The 2021–2022 wet season brought roughly 602 mm of precipitation, which is relatively generous given a historical mean of 473 ± 198 mm.

16 trees were selected from inner rows in both 10-year-old Navel orange and 15-year-old Shelly mango orchards. Trees from each species were selected based on a superficial and internal evaluation of health. Tree Core samples were taken and assessed for disease, and trees with multiple stems, as well as orange and mango trees with diameters less than 10 and 30 cm, respectively, were excluded. No other experiments have been performed on the trees that participated in out study (and generally the entire orchard surrounding them); these orchards were mainly a source of income for the farm. Navel orange trees were spaced 4×4 m, while Shelly mango were spaced 6×6 m, and each receives drip irrigation from mid-spring to late autumn (flow rate of 1.6 L h^−1^), according to guidelines from the Ministry of Agriculture (typically divided into weekly doses of 2-3 irrigation days). Soil moisture content was never lower than guidelines during the study period, and no deficit irrigation was applied. Soil in both orchards was a loamy Hamra, a red soil inherently rich in iron, with good drainage. The mango orchard has also received tree clipping compost for a decade, which has created a dark top layer. Both orchards were on an incline of approximately 2% eastward.

### 2.2 Tree and soil sensors

In each orchard, East-30 sap flow sensors (Pullman, WA, USA) capturing 2-directional flow were placed in the stem of 16 trees, at roughly 30 cm above the ground and before branching, and eight trees were fitted with a modified Teros-12 soil sensor (Meter Group, Pullman, WA, USA) to measure stem water content (StWC, method by Matheny et al. 2017). Nine soil moisture sensors (Acclima TDR-315H, Meridian, ID, USA) were installed in each orchard, where three sensors in three rows measured soil moisture at three depths. Based on an initial excavation, orange root density was similar to depths reported by Wagner et al. (2021) and Cohen and Cohen (1983), with high density in the upper 30 cm layer. Therefore, the three chosen depths in the orange orchard were 20, 40, and 80 cm. Mango, alternatively, has fewer superficial fine roots and a deep taproot with some lateral roots. Our findings were consistent with Santos et al. (2014), where root density was spread between 30 cm to 90 cm depth, and therefore the three chosen depths were 30, 60, and 90 cm. Due to challenges in excavation with a Little Beaver motorized auger, orange and mango soil sensors were placed 160 cm and 140 cm, respectively, away from the stem of the nearest tree; despite the distance, both were installed under the crown and within the root network of the tree. Soil water content was monitored to ensure no soil water deficits occurred (at all depths). Data was collected from all sensors every two minutes and was batched and transferred wirelessly to our server every 30 minutes.

### 2.3 Meteorological data

A meteorological station was placed on a 3-m tall tower in the navel orange orchard, with sensors measuring air temperature, relative humidity, incoming and outgoing short- and long-wave radiation, and photosynthetically active radiation. A smaller, 2-m tall, tower was placed in the adjacent mango orchard, measuring air temperature and relative humidity. For historical data, sets of 3-hourly and daily temperature and relative humidity data was available from a nearby Israel Meteorological Service station in Beit Dagan (station 2520, 10 km distance, IMS (2024)), with minor gaps filled with data from a second nearby station in the same city (2523). 70-year historical daily precipitation data was also collected from station 2520. Daily precipitation data for October 2021 to January 2022 was obtained from station 2520, and from January to April 2022, sub-hourly precipitation data was available from a monitoring station within the farm.

### 2.4 Data analysis

#### 2.4.1 Calculating VPD percentile thresholds

As the first set of data processing, extreme VPD events had to be characterized as events that exceeded historically, statistically high VPD values. From 3-hourly historical temperature and humidity data, we calculated VPD using the Pynotes in Agriscience VPD function (Patrignani, 2020). The 3-hourly VPD values were then aggregated into daily medians across all 56 years of available data (1964-2020). We preferred daily medians over daily averages because VPD is not normally distributed. We then grouped all values according to day of year (irrespective of year), and applied a 14-day rolling window. Finally, we computed the following VPD percentiles inside this window: 50th (median), 80th, 85th, 90th, and 95th. Figure 2a shows historic monthly rainfall averages and the computed historic percentiles. Panel b focuses on the 2022 spring season, showing the daily maximum VPD overlaying the historic percentiles, while panel c shows a heatmap with hourly VPD values for the same period.

**Figure 2:**
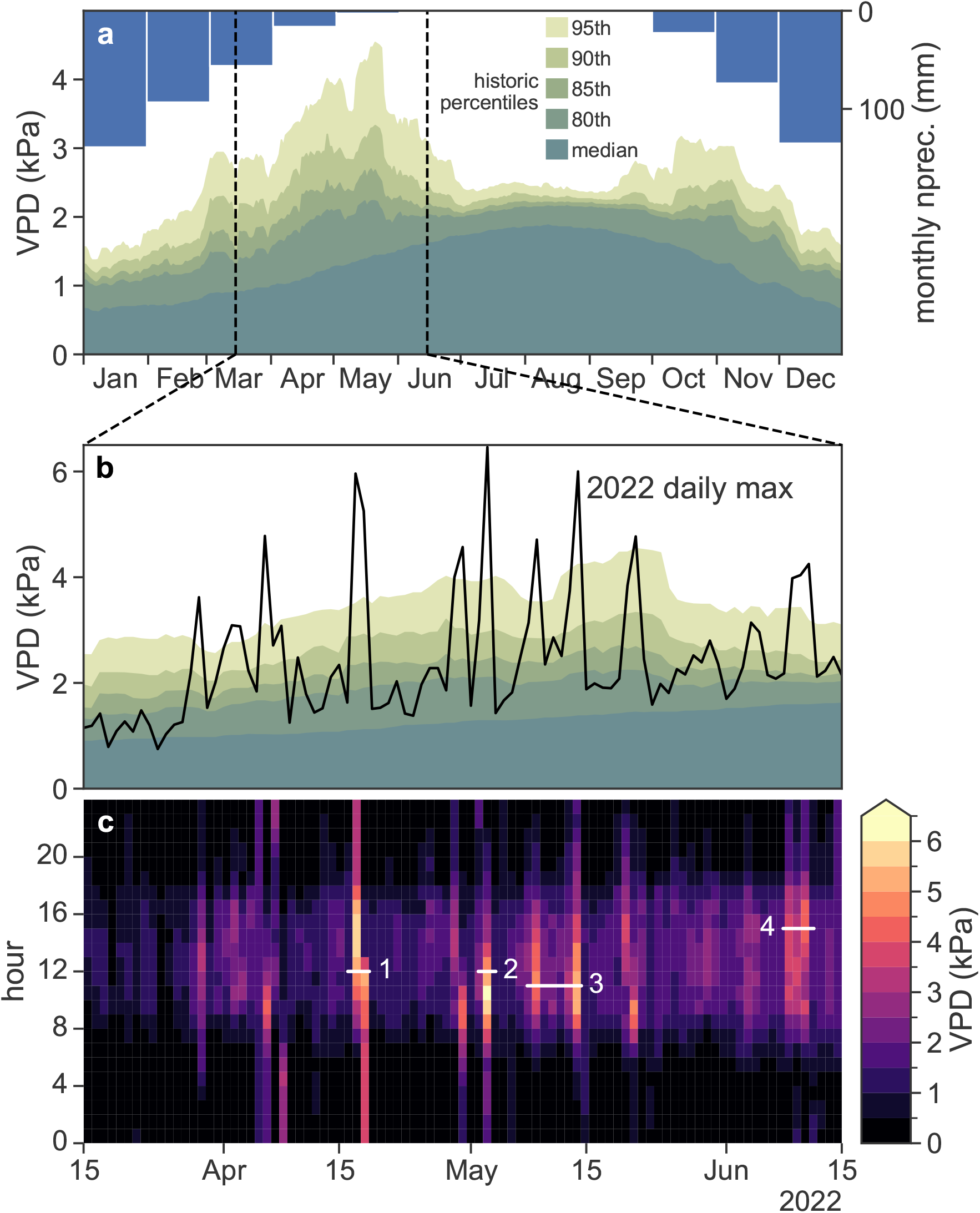
VPD is widely variable in spring and fall (IQR of 0.65 kPa and 0.59 kPa, respectively), while median VPD is greatest over summer, but with less than half the spread (IQR= of 0.28 kPa). Panel a: historic monthly precipitation averages (blue) indicate the high seasonality of the mediterranean climate, and historic VPD percentiles (green shades) suggest a bimodal pattern of VPD variability. Panel b: the daily maximum VPD (black) for the 2022 spring period is shown overlaying the historic percentiles. Panel c: hourly VPD for this same period is shown as a heatmap.

#### 2.4.2 Smoothing

We used SciPy’s Savitzky-Golay filter (Virtanen et al., 2020) to perform smoothing of all time series. While smoothing with regular running averages has the effect of spreading the signal (lowering peaks and raising valleys), the polynomial best-fit used by the Savitzky-Golay filter can capture nuanced dynamics in the data, such as sharp changes in the time series dynamics.

#### 2.4.3 Daily trend and its derivative

We used Python’s statsmodels package (Seabold and Perktold, 2010) seasonal decompose tool to split StWC into three components: trend, seasonality, and residuals. The goal was to get the daily trend dynamics of StWC without the expected daily fluctuations (seasonality) and jitter (residuals). The trend is calculated by the algorithm with a running average whose window size is one day, yielding the net change in StWC unrelated to tree buffering activity. In order to get the rate of change (derivative) of this trend, we used the Savitzky-Golay filter with a best-fit 2^nd^ degree polynomial, corresponding to a linear rate of change over the running window used by the filter.

## 3 Theory and calculation

### 3.1 Quantifying the Stem Water Storage and the Cumulative Heat Unit

To obtain the mass of StWS [kg], first the sapwood area of an individual tree (*A*_sw_ [cm^2^]) was calculated. Assuming circular cross sections, the sapwood area is the area of a ring whose outer radius is *D/*2 and its inner radius *D/*2 − *h*, where *D* is the stem diameter [cm] and *h* is the sapwood thickness [cm], yielding

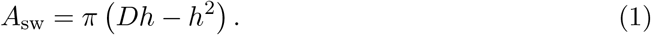

The thickness is the mean thickness of active xylem by tree type, as determined from core samples of each tree. Thickness was approximated, based on core samples, to 2 cm in orange trees and 6 cm in mango trees.

To calculate StWS [g], StWC [g cm^−3^] was multiplied by the tree’s sapwood volume, yielding

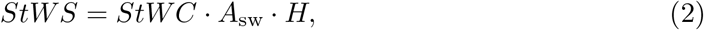

where *H* [cm] is the stem height. The Cumulative Heatwave Unit (CHU, [kPa h]) for a single event was calculated as

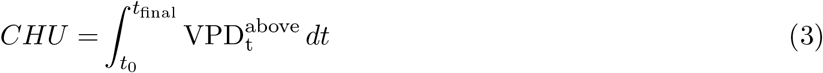

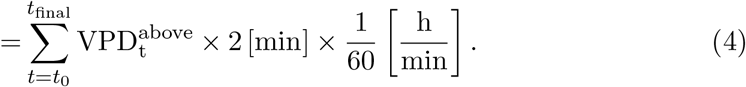

The integral expression is the general definition of CHU, giving the area of VPD above a given threshold 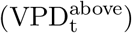 between *t*_0_ and *t*_final_. The second line shows the discrete expression we applied to our time series, where all measurements come in a 2-minute frequency, and the factor 1/60 converts minutes to hours.

### 3.2 Derivation of relationship between StWS and CHU

We start with the common relationship between transpiration *E* [mmol m^−2^ s^−1^] and VPD [kPa],

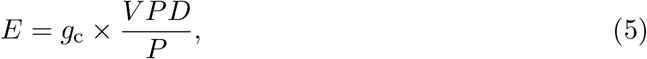

where *g*_*c*_ [mmol m^−2^ s^−1^] is the canopy conductance, and *P* [kPa] is atmospheric pressure. *g*_*c*_ can be be approximated to vary linearly (slope provided by coefficient, *a*_1_) with whole-plant leaf-specific hydraulic conductivity *K*_*L*_ [mmol m^−2^ s^−1^ MPa^−1^] (Hubbard et al., 2001) as

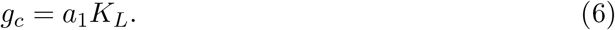

By substituting Eq. (6) into (5), integrating over time, expanding the volume of transpired water into a linear combination of water donated from storage pools (mainly stem and roots), and linking the use of these pools to various VPD conditions, we were able to obtain the following relationship for a change in StWS driven by statistically high VPD (VPD above percentile threshold):

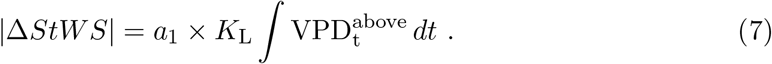

We show a the full derivation in section S1 of Supplementary Material.

The magnitude of the heatwave can be quantified by the final term in the previous equation, identified in Eq. (3) as CHU, and it incorporates both the intensity and duration of the extreme VPD event. Finally, we find the relationship between the absolute changes in stem water content and the heatwave magnitude:

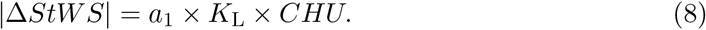

### 3.3 Defining a heatwave

To define a dry heatwave in relation to a tree, VPD above high percentile thresholds was related to the trend change, and rate of change of the trend, of StWC in each species. Similarly, statistical based definition of a heatwave (not necessarily dry or wet) could be based on the air temperature rather than the VPD, but that was not used in this study. Figure 3 shows the VPD above these percentiles (panel a) and the dynamics of trend StWC and its derivative, for both orange and mango trees (panels b and c), over a 3-month period. Both species experience reduction in trend StWC (thick solid blue lines) when VPD is at the 80th percentile or higher. When the trend in StWC declines, its rate of change is negative (thin gray lines). Trend in StWC has a delayed recharge (tied to physical conductivity (Ewers and Cruziat, 1990) and the lag between sap flow and transpiration (Preisler et al., 2022), however, the negative rate quickly returns to zero or becomes positive once the event is over. From this common characteristic occurrence in both species, heatwave events were thus defined as the periods at which both VPD was above the percentile thresholds and when the derivative was simultaneously negative.

**Figure 3:**
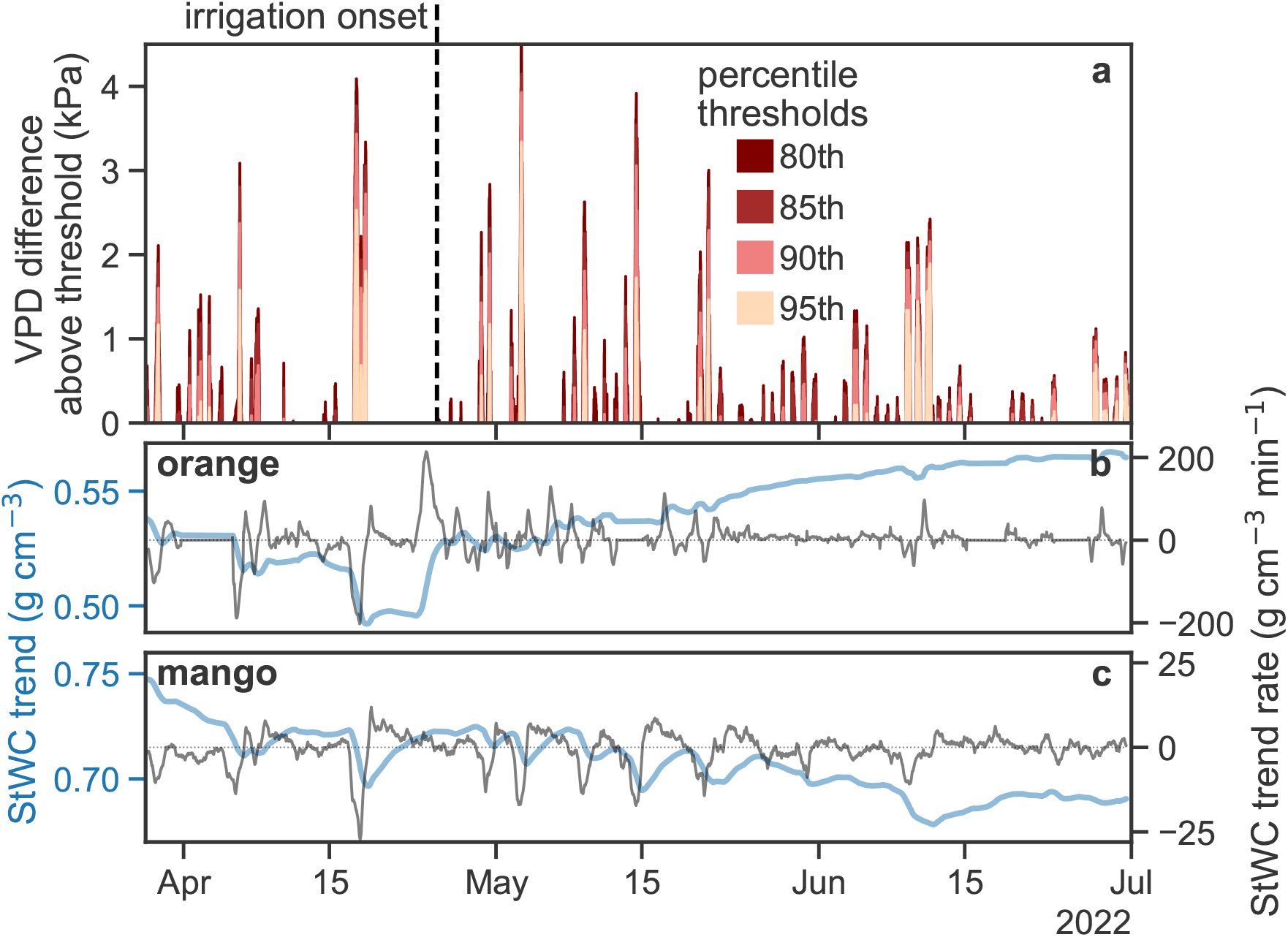
VPD above high percentile thresholds causes reductions in trend StWC in both species of trees, in particular in Navel orange trees in the rainfall period. Panel a: Δ*V PD* above percentile thresholds (80th, 85th, 90th, and 95th). Panels b and c: trend StWC [g cm^−3^] and its derivative [g cm^−3^ min^−1^] for Navel orange and Shelly mango trees.

The total reduction in StWC was calculated as the difference between the maximum and minimum StWC in an event (similar to the maximum daily shrinkage described by Dietrich et al. (2018), but for the trend StWC for a respective event, which can be multiple days). It was found that there is an important change in stem water management between the rainfall and the irrigation period in both trees (discussed in results), which brought us to assess the relationship between StWS and CHU separately in these distinct periods.

There are incidents where there is a negative trend rate with no corresponding event, which may be explained as instances when StWC use is not related to transpiration (Preisler et al., 2022), or that transpiration demand was relatively—but not significantly— higher than a typical diurnal flux and was therefore carried forward during the detrending step. The change in StWC trend in these instances is negligible.

## 4 Results

### 4.1 Climate and variability

The 2021 wet season (October 2021–April 2022) received 602 mm of precipitation, which stands within historical norms (474 ± 198 mm). Within these winter months, air temperature was also characteristic for the region, with few hotter-than-average days. Within the study period, though there were numerous days with exceedingly high temperature and VPD (see Fig. 1 and 2c), none were record-breaking. Overall, the mean climate during the study period and in the preceding wet season can be considered standard for the region.

We performed a preliminary check for heatwaves in the study period by applying the variety of definitions given in Table 1. It should be noted that, when we included statistical variability as one standard deviation for air temperature, none of the definitions produced any heatwave events. If variability is excluded, as it is in each definition, then the resulting heatwaves were identified and are shown in Table 1, where applicable.

**Table 1:** Respective heatwaves in study period according to existing definitions.

| Source | Heatwave definition | Dates (DD/MM) in 2022 study period |
| --- | --- | --- |
| Schär et al. (2004) | Mean summer (June–August) temperature $\geq 3^\circ\text{C}$ of historical mean. | None. [historical: $25.5^\circ\text{C}$, 2022: $27.5^\circ\text{C}$] |
| Sun et al. (2014) | Daily maximum temperature $\geq 35^\circ\text{C}$. | 17–18/04, 03/05, 14/05, 21/05, 11/06 |
| World Meteorological Organization (WMO) (2022) | At least five days with a sustained daily maximum temperature $> 5^\circ\text{C}$ of the historical daily maximum temperature. | None. Longest event only three days. |
| Cochavi et al. (2021) | Beginning: daily VPD average $> 0.8\text{ kPa}$ of previous day. End: daily VPD average $> 1.0\text{ kPa}$ and daily temperature average $> 5^\circ\text{C}$ of following day. | 28/03–01/04, 17–19/04, 02–04/05, 09–10/05, 20–22/05, 09–12/06 |

Heatwave definitions that span the variety of formulations, and their occurrences during the study period, are shown in Fig. 1. The definitions by Schär et al. (2004), Sun et al. (2014), and Cochavi et al. (2021) were developed for Europe, China, and the Mediterranean, respectively, whereas the definition by the WMO is a global definition.

To quantify a dry heatwave in CHU, first, the magnitude of daily variability in VPD was calculated for the four high percentile thresholds: the 80th, 85th, 90th, and 95th percentile. The results are plotted in Fig. 2a, where we see that variability in VPD is highest in spring (interquartile range, IQR=0.65 kPa) while the median is relatively low, but rising. In summer, median daily VPD is greater than throughout the rest of the year (*>* 1.5 kPa daily), however there is far less variability (IQR=0.28 kPa). In other words, VPD in spring is more than twice as scattered as the VPD in summer, which, historically, has a record peak of 3.88 kPa versus 6.71 kPa in spring. In the Eastern Mediterranean climate, this is due to the combination of consistently high (and converging) daily maximum and minimum temperatures in summer, as well as the relatively constant daily humidity, which produces a narrow range of summer daily VPD values. This pattern in VPD percentile scatter is similar to the temporal pattern of recorded hamsin events in the same region found by Tatarinov et al. (2016).

### 4.2 VPD at a finer timescale reveals species sensitivity

The intensity and duration of VPD at a fine scale (sub-daily) was found to be highly relevant for describing a dry heatwave, as opposed to the common daily maximum methods. Figure 2 shows VPD values between 15 March and 15 June, 2022, where 2b gives daily VPD maxima and 2c gives a heatmap of hourly VPD averages. Maximum daily VPD reaches numerous peaks above 4 kPa, signifying values greater than the 95th percentile in spring. It was found that the intensity and duration of a statistically rare VPD event can vary greatly both between events of a same percentile (intra-percentile events), and events of different percentiles (inter-percentile events). For example, the most intense VPD event in the study period (event 2 in Fig. 2c) reached a maximum daily VPD of 6.46 kPa and remained above the 95th percentile for 6 hours, while event 1 reached a lower peak VPD of 5.96 kPa, but remained above the 95th percentile for 12 hours, with significantly high nighttime VPD. Both events 1 and 2 contrast in comparison to event 4, which remained above the 95th percentile for 33 hours in a 3-day event, but only reached a peak VPD of 4.10 kPa. Each of these events (1, 2, and 4), however, can be categorized together as intra-percentile events due to the shared rarity of their VPD intensity (95th percentile).

In comparing inter-percentile events, we found that the duration of VPD intensity become even more significant as events in lower percentiles tended to persist for longer periods of time. Event 3, which reached a maximum VPD of 5.43 kPa (though only remained above the 95th percentile for 6 hours on two non-consecutive days), surpassed the 80th percentile for 71 hours over 8 days. Since intensity and duration of a climate driver commonly carry the same weight in heatwave definitions (total weight split 50/50), it is evident from this analysis that sub-daily percentile-based quantification improves the characterization of events by capturing the changing dynamics between these factors both across intra- and inter-percentile events. Moreover, since there are notable differences in intensity and duration between inter-percentile events, there is room to assess at which percentile threshold the relative events have the most significant effect on the tree, by species.

### 4.3 Change in StWS and CHU, and net change in StWC

CHU and |ΔStWS| were calculated according to subsections 3.1 and 3.3, respectively, for CHU at three percentile thresholds (80th, 90th, and 95th — the 80th and 85th percentile were very similar) and the results are plotted in Fig. 5. Some notes about the dynamics of the variable, CHU: CHU does not tend to increase by percentile threshold, which can be explained by 1) 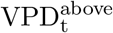 (section 3.1) and the measured VPD, itself, are not necessarily coupled, and that 2) VPD events of a higher percentile threshold tend to be shorter in duration than events of a lower intensity (see Fig. 2c). CHU magnitude is also species-dependent, since it is based on the co-occurrence of a high VPD with a negative StWC trend rate (which is species-dependent). Due to the latter, the number of events (n) is also species-dependent, and does not necessarily increase or decrease according to percentile. This can be seen, for example, in mango trees, where the most events in the irrigated period occur at the 90th percentile (n=13). There was also no significant difference found between the dry heatwave events of the 80th and 90th percentiles (*p* = 0.277), while the response by season and between species was found to be significant (*p* = 0.051 and *p <* 10^−3^, respectively), where the slope of the StWS response to dry heatwaves in orange trees was much lower than the response in mango trees (0.003 and 0.115, respectively; statistics are summarized in S4). Therefore, the best relationship between |ΔStWS| and CHU for a given species and period was decided to be that with the greatest significance according to R^2^ and n. For each relationship, the level of agreement between ΔVPD and negative StWC trend rate (by section 3.3), ‘ag’, is also given (explained in section S2 of Supplementary Material).

**Figure 4:**
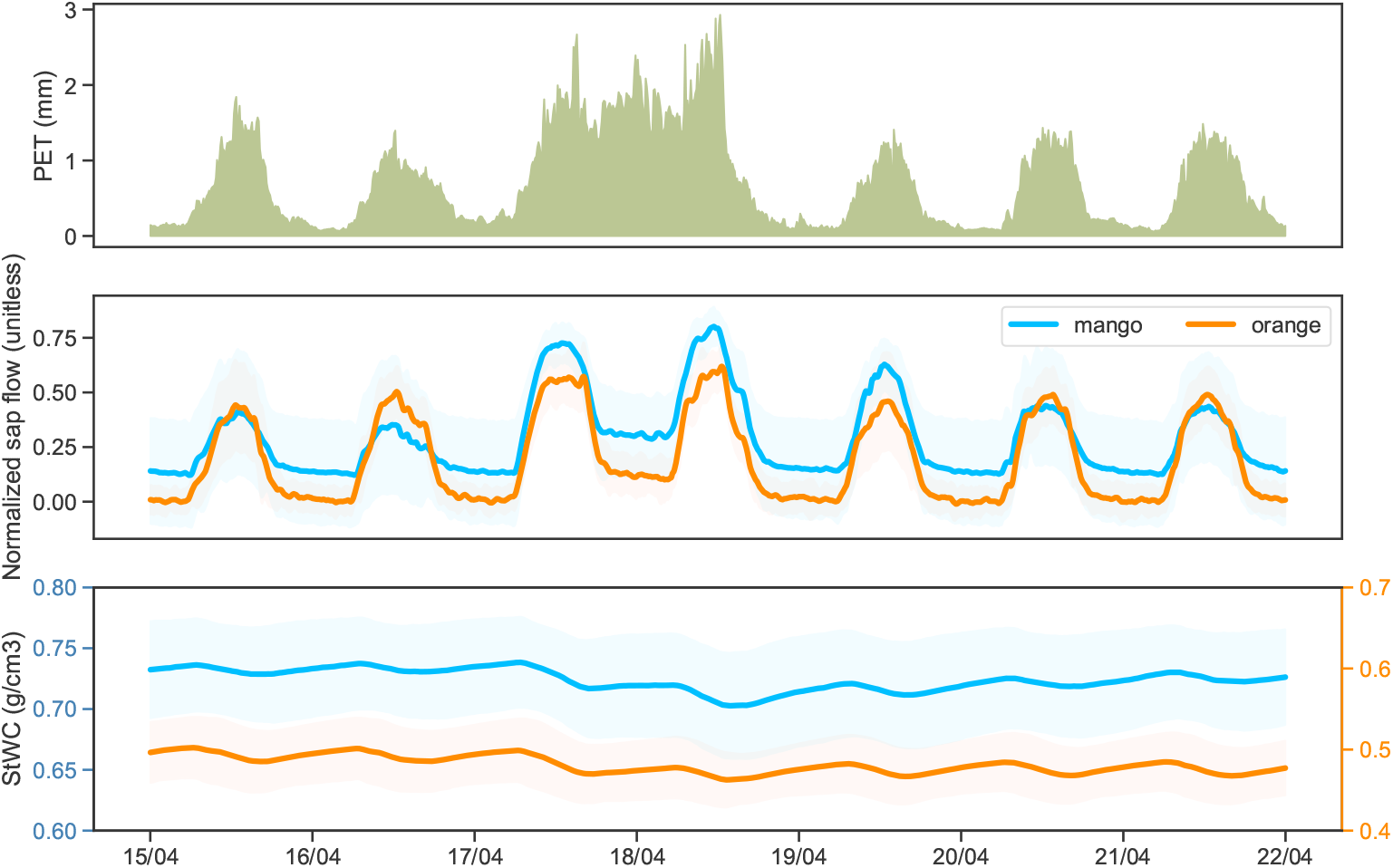
Direct response in sap flow and StWC to increased transpiration demand. Sap flow sharply increases, especially overnight, in both species during high transpiration demand (April 17th–18th; estimate from Penman-Monteith (Vremec and Collenteur, 2021)), though reaches a midday plateau in orange trees, while StWC reduces over this period without full recovery in the following days.

**Figure 5:**
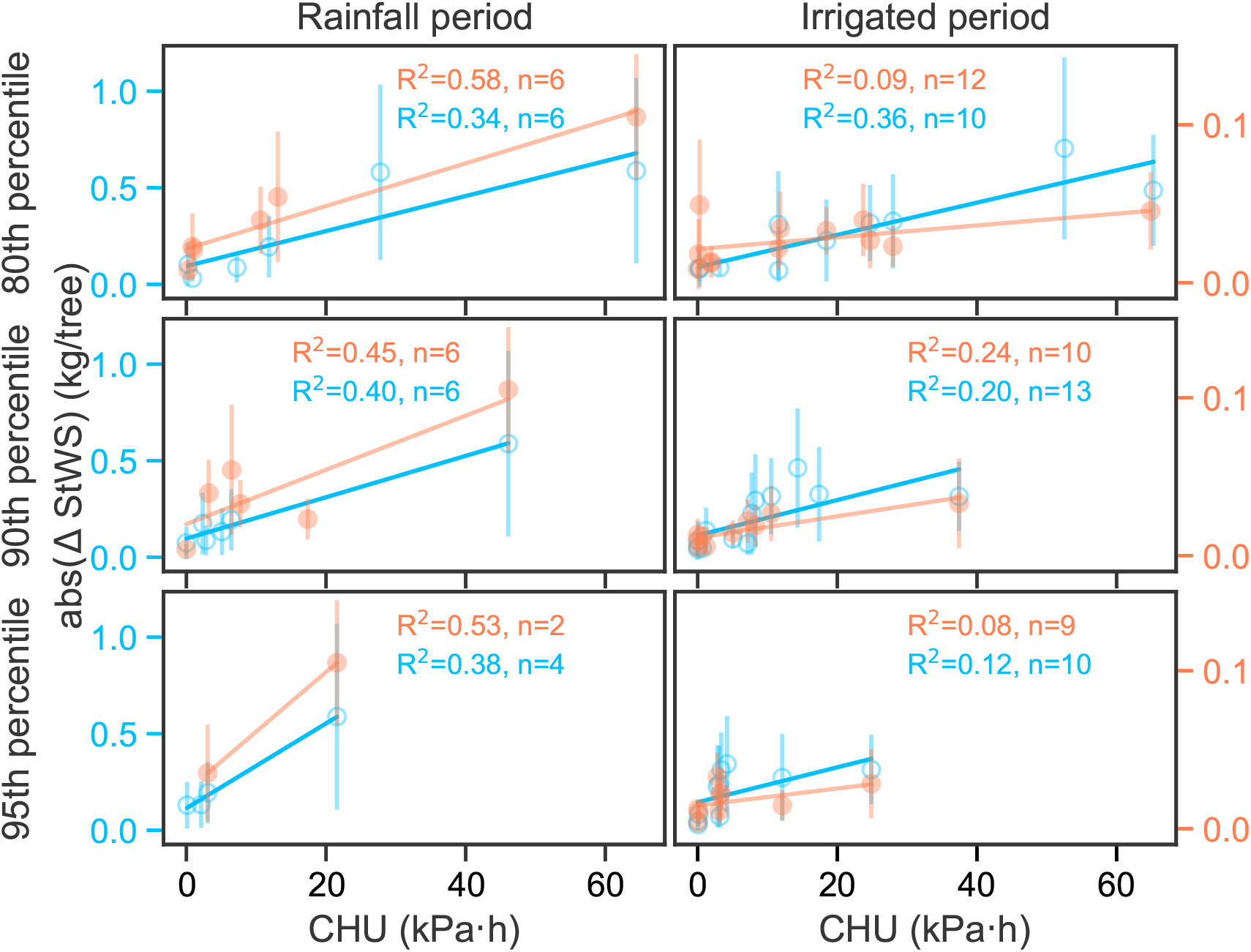
Change in StWS [kg tree^−1^] by CHU [kPah] in orange and mango trees (colored orange and blue, respectively). Plots are organized in columns according to rainfall and irrigated period, and in rows by percentile thresholds (80th, 90th, and 95th).

In orange trees in the rain-fed period, the best-fit relationship between |ΔStWS| and CHU is at the 80th percentile (R^2^=0.58, n=6; ag=64.7 %), whereas no statistically significant relationship exists in the irrigated period (R^2^ *<* 0.25). |ΔStWS| and CHU relationship in mango trees is best described by events at the 90th percentile (R^2^=0.40, n=6; ag=98.5 %) in the rain-fed period, and at the 80th percentile (R^2^=0.36, n=10; ag=88.7 %) in the irrigated period. We believe that the statistical significance of the VPD events by percentile should improve in experiments with longer study periods, where more events can be captured by season, since the relatively low number of events and input variables to the simple regression and mixed linear model likely affected the output.

Regarding change in StWS as a result of heatwaves, there was a peak mean loss in StWS of 105 g (STD=36.7 g, CHU=64.4 kPah) per orange tree on April 17th-18th. This amount is reasonable, considering the peak total daily uptake of water reported by Mishra et al. (2021) for 5 and 15 year-old orange trees to be 1.1 to 5.0 kg day^−1^, respectively, given that more than 95% of this uptake is lost daily to transpiration (McElrone et al., 2013). For a 10 year-old tree, the remaining portion of uptake, which normally contributes to cell expansion and growth (McElrone et al., 2013), would equate with the given parameters to a mean 150 g day^−1^. In mango trees, the same event in the rainfall period (April 17th–18th) resulted in a mean loss of 588 g (STD=439 g; CHU=46.1 kPa h), however the period-peak mean loss of 703 g tree^−1^ (STD=431 g; CHU=52.5 kPa h) occurred in the irrigation period. These amounts are comparable to an estimated 830 g day^−1^ of water up-taken but not transpired in 11 year-old Palmer mango (which has a similar sapwood area to Shelly mango), based on the measured 16.6 kg day^−1^ of water transported in sap flow on bright sunny days (Oguntunde et al., 2011). Given these peak losses, it appears that a heatwave of roughly 50–65 kPa h will cost nearly the entire daily water uptake of a well-watered tree, based on the relative percentile in orange and mango trees.

We found that this price — nearly the entire daily water uptake being transpired, leaving little to no water for stem storage replenishment — varied by species. Orange trees, which began in the study period with a peak StWC of 0.55 g cm^−3^, were most sensitive to heatwaves in the early spring, during which time StWS was not immediately replenished after a heatwave (Fig. 4). This changed in May, as daily temperatures continued to rise and likely regulated stomatal conductance and transpiration, resulting in a portion of the daily uptake going to replenish StWS, as by the end of the study period StWC even slightly exceeded the initial density (reaching 0.57 g cm^−3^, Fig. 3). In mango trees, however, the trend in StWC decreased over the study period from 0.75 g cm^−3^ to 0.69 g cm^−3^ (Fig. 3). These differing responses between species can also be seen in measurements of their respective diurnal sap flow during a heatwave, as seen in Fig. 3. Sap flow in orange trees reached a midday plateau on the April 17th heatwave (likely associated to a reduction in stomatal conductance (Ortuno et al., 2006)), while, oppositely, sap flow in mango trees increased to a midday peak (likely due to continued transpiration (Lu and Chacko, 1996)). Finally, it was also found that the more negative StWC trend rate in orange than in mango trees (Fig.3) is consistent with the more negative “depletion rate” in isohydric than anisohydric species (Dietrich et al., 2018).

## 5 Discussion

### 5.1 Change in StWS in a heatwave

In the Mediterranean climate, StWS is known to help buffer morning to midday transpiration demand in the time before water uptake from the soil can reach the canopy (Preisler et al., 2022). Here, we have demonstrated that beyond this diurnal flux, StWS further supports the additional transpiration demand during a heatwave (see Fig. 3). This additional loss was found to be up to 105 g (STD=36.7 g) and 703 g (STD=431 g) per event and tree in orange and mango trees, respectively, which is similar in value to the estimated remaining daily water uptake from the soil that is not transpired for each respective species of 150 g and 830 g (McElrone et al., 2013) for similar cultivars (Oguntunde et al., 2011; Mishra et al., 2021). This suggests that there would be little to no stem water recharge during a heatwave, which is what we observed from our StWC measurements (see Fig. 4), along with relatively high sap flow during the events.

To further explore the value of the StWC measurements, which are obtained from a relatively recent technique in which a modified soil sensor is installed directly within the sapwood area in the stem (Matheny et al., 2017), we estimated two values from our data to compare to literature values. First, we compared the theoretical change in stem diameter due to water volume reduction to the inferred losses by diameter changes in tree water deficit (ΔW) (Zweifel et al., 2005) and maximum daily shrinkage (MDS) (Dietrich et al., 2018) of species with similar water management strategies in heat and high-VPD exposure. Given the diameter and height of each study tree’s stem, as well as its respective volumetric change in StWC in a heatwave, we calculated a ballpark estimate for the theoretical mean peak change in stem diameter caused by a heatwave, resulting in 0.53 and 0.96 mm in mango and orange trees, respectively. Broadly comparing these values to studies on other anisohydric and isohydric species, respectively, we found that these values were greater than the shrinkage in trees in non-heatwave conditions (MDS 0.075 and 0.2 mm, respectively, derived from Hahn and García (2022); García-Tejero et al. (2012)), and more comparable to trees experiencing heat-treatment or high VPD (ΔW of 0.25–1.0 and 0.3–0.4 mm for the measured anisohydric and isohydric species, respectively (Zweifel et al., 2005; Ruehr et al., 2016)). Though our calculated changes in diameter are ballpark estimates, the magnitudes compare reasonably well (a MDS difference of less than 1 mm) to species with similar water management strategies, especially when under heat and VPD stress.

Second, we sought to estimate and compare the values obtained for stem hydraulic conductivity (K_stem_) for each species, given the relationship derived in Eq. (8) between ΔStWS and CHU. According to it, K_stem_ should be proportional to the slopes of the regression curves (Fig. 5) by a factor *a*_1_. As *a*_1_ has been demonstrated to be less than 10 (ranging within a single order of magnitude) in trees with both isohydric and anisohydric water management (Hubbard et al., 2001), we compared the magnitudes, log_10_K_stem_, derived from our results with those of literature values (see Table 3 in S3). Notably, we found that log_10_K_stem_ in Shelly mango did not change between the rain and irrigated seasons and ranged between 0.92–1.34, which aligns well with the calculated log of K_stem_ of mango cultivars from the literature, which range between 0.69–1.16 (Normand et al., 2008). In Navel orange, log_10_K_stem_ was only positive in the rain season, ranging from 0.13–0.58, while the range of a similar cultivar in the same region was found to be greater (1.08–1.15) (Cohen et al., 1987). The estimated log_10_K_stem_ for both cultivars in this study were less than an order of magnitude different from the range of calculated log_10_K_stem_ for similar respective cultivars in the literature, which suggests that the relationship derived in subsection 3.2 relating ΔStWS and CHU is accurate. Further details on this analysis can be found in S3.

**Table 2:** Events that should have been considered heatwaves according to ΔStWS_min_ by the events outlined by each definition, relative to their best-matched percentile threshold.

| | Heatwave dates in<br>2022 study period<br>(DD/MM) | Percentile<br>threshold of<br>defined<br>events, relative<br>to species:<br>Navel orange &<br>Shelly Mango | $\Delta\text{StWS}_{\min}$ [g] of<br>the given heatwave<br>dates for<br>Navel orange &<br>Shelly Mango | Events in percentile<br>with<br>$\Delta\text{StWS} >$<br>$\Delta\text{StWS}_{\min}$<br>(DD/MM) |
| --- | --- | --- | --- | --- |
| Sun et al. 2014 | 17–18/04, 03/05,<br>14/05, 21/05,<br>11/06 | 95th (both) | 15 & 273 | Orange: 29/03, 29–<br>30/04, 27–30/06<br>Mango: NA |
| Cochavi et al. 2021 | 28/03–01/04, 17–<br>19/04, 02–04/05,<br>09–10/05,<br>20–22/05, 09–12/06 | 80th (both) | 23 & 194 | Orange: 07–08/04,<br>25–27/04,<br>29–30/04, 24–31/05<br>Mango: 29–30/04,<br>26–31/05 |

**Table 3:**
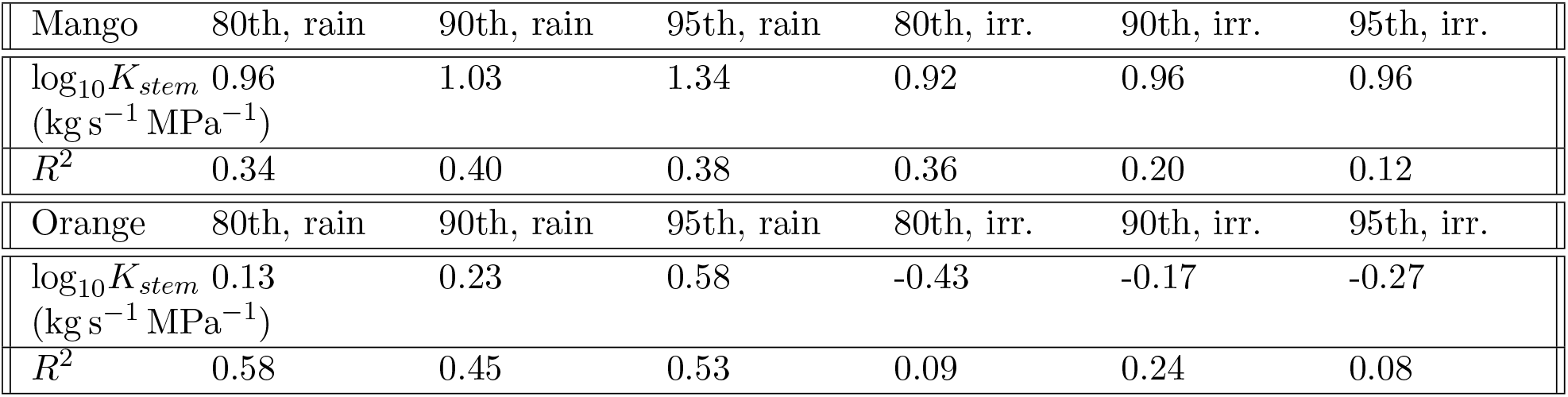
The magnitude of *K*_*stem*_ in mango relative to percentile threshold and period. The magnitude is calculated from the slope of the regression curves in figure 5 (log(slope/a1)).

### 5.2 Heatwave definition

Given the complex interplay between VPD intensity and duration outlined in section 4.2, we evaluated the definitions from Table 1 (those that produced heatwave dates) on their ability to determine significant events that affect StWS in the trees. In comparing the dates obtained from each definition in Table 1 to those obtained by applying the method in section 3.3, the dates from Sun et al. (2014) correspond well to events in the 95th percentile for both species, while the dates obtained from Cochavi et al. (2021) match best with those within the 80th percentile (also for both species). Note that all events defined by respective authors were also obtained by our method, however, for each species’ percentile and season, our method found additional events. This brought us to qualify the significance of the definitions’ heatwaves according to loss in StWS. Table 2 shows the minimum change in StWS (ΔStWS_min_) that was lost per species according to each respective heatwave definition dates, and gives the subsequent dates that should have been considered heatwaves, given that the tree’s reduction in StWS was greater than ΔStWS_min_.

According to the dates that were not identified by Sun et al. (2014), but had an equal or more significant effect on StWS, the definition does not include three events that specifically affect Navel orange trees. Two of these events saw daily maximum temperatures reach 33.8 ^*°*^C and 32.6 ^*°*^C (29–30/04 & 27–30/06, respectively); the third had a milder maximum (25 ^*°*^C), but occurred earlier in the season (29/03). The first two events show that the application of a maximum daily threshold for temperature can exclude events that are equal or more significant to tree WS as those above the threshold (likely due to the duration component). The third event highlights the value of a definition based on statistical variability in a seasonal climate, as temperature (and VPD) exceeds the 95th percentile in early spring at a lower value than in late spring and summer (see Fig. 2a). Given our definition of a heatwave, another way to describe this is that the former definition misses events at the 95th percentile with a CHU of 2.80–3.20 kPa h.

As the heatwaves found using the Sun et al. (2014) definition are best described by the 95th percentile, and StWS in orange and mango was found to be sensitive to heatwaves at at least the 80th and 90th percentiles, respectively, this definition under-estimates the total number of events by three events in orange and twelve events in mango. The definition from Cochavi et al. (2021) better agrees with our findings, given that the dates correspond to the 80th percentile — the same percentile that best describes Navel orange and Shelly mango in the rainfall and irrigation period, respectively. It did, however, miss four events in orange and two in mango. The dates left unidentified commonly meet the criteria for ΔVPD between day_final_ − day_final+1_ *>* 1 kPa, however the temperature differences were not greater than 5 ^*°*^C. Moreover, the events are characterized by relatively low peak VPD (less than 3.62 kPa), but lasting an average of 3.8 days. Essentially, it is seen again that a heatwave definition built binarily upon thresholds will inevitably be unable to capture to wide variety of events due to the interplay of intensity, duration, and relative statistical rarity (relative to the season). In terms of CHU, this definition misses events at the 80th percentile ranging between 10.30–18.40 kPa h.

It is intuitive and also clear from CHU plotted in Fig. 5 that heatwaves (and the resulting impact on tree WS) continually increase in magnitude due to increasing VPD intensity and duration. We found, however, that definitions that do not incorporate the nuanced interplay between intensity and duration on a continuous scale actually miss some meaningful occurrences of these events within the overall range. Moreover, when statistical rarity is incorporated in these definitions (as 1 STD for each climate variable), no heatwave events were recorded, despite VPD reaching above 5 kPa in numerous events. We found that our framework for identifying heatwave events from the tree’s perspective succeeded in relating statistically rare VPD events with trend changes in StWS principally by working on a finer (sub-daily) timescale, within which we could accurately combine both VPD intensity and duration. It should be noted that this framework relies on historical data for the studied region for computing percentiles, however no fixed thresholds are ever imposed (as is the case in Schär et al. (2004); Urban et al. (2017); Drake et al. (2018); Sun et al. (2014); Cochavi et al. (2021)). As such, the framework for computing heatwave magnitudes can be widely employed in studies across geographic locations, since there is no component related to a single geographic location or climate condition. As far as we have found, ours is the only definition of a heatwave that offers a magnitude for events, with units, that is calculated from statistically high VPD and duration, and which, explained by theory and practice, is defined according to a significant response from the relative tree species that experiences it.

## 6 Conclusion

In this work, we provided a definition for the magnitude of a heatwave based on its statistical rarity as a VPD event integrated over its duration, and the significance of the tree’s response in terms of StWS. We found that there is a statistically significant and opposing response between isohydric Navel orange and anisohydric Shelly mango trees, where mango trees lose a significantly greater factor of StWS than orange trees during a heatwave (0.115 versus 0.003, respectively), and overall see a decline in StWC between the wet and the irrigated dry season (from 0.75 to 0.69 g cm^−3^) while StWC in orange trees slightly increases (from 0.55 to 0.57 g cm^−3^). We found that mango trees increased in sensitivity to heatwaves, while orange trees demonstrated resilience to StWS losses between the wet and irrigated dry season. By quantifying a heatwave, we were also able to compare the magnitudes of heatwave events and show that both the duration and intensity are significant factors, and, importantly — that the intensity of a heatwave is relative to the species experiencing it. The relationship developed here between the additional loss in StWS during a heatwave can be used to predict, for example, additional irrigation demand in an orchard during a heatwave, and can be employed in any geographic region with a reasonable historical climate-record.

## Abbreviations

CHU: Cumulative heatwave unit (kPa h)
EWE: Extreme weather event
IQR: Interquartile range
MDS: Maximum daily shrinkage
STD: Standard deviation
StWC: Stem water content (g cm^−3^)
StWS: Stem water storage (kg)
WS: Water storage
VPD: Vapour-pressure deficit (kPa)

## Acknowledgements

We would like to thank the following researchers, technicians, staff and outside consultants for their unique contributions to the quality of the research: Dr. Ilana Shtein, Ariel University, for her guidance in obtaining and analyzing core samples; Dr. Ashley Matheny, University of Texas at Austin, for assistance in applying her published method for measuring StWC; Gil Lerner, the Hebrew University of Jerusalem, for his assistance in assembling the climate towers and in preparing the Teros sensors; the consultants from EnviroManager, who were readily available for assistance with datalogger and tower communication issues; and the many farm staff who monitored and quickly repaired the punctures to irrigation from thirsty birds, and maintained the Ministry’s prescribed protocol.

## Author contributions

**Laura Rez**: Conceptualization, data curation, formal analysis, methodology, visualization, writing (original draft) **Justine Missik**: Software, writing (review and editing) **Gil Bohrer**: Conceptualization, funding acquisition, methodology, supervision, writing (review and editing) **Yair Mau**: Conceptualization, funding acquisition, investigation, methodology, project administration, supervision, visualization, writing (original draft).

## Funding sources

This research was supported by Research Grant Award No. IS-5304-20 from BARD, The United States-Israel Binational Agricultural Research and Development Fund.

## Appendix

### A1: log_10_K_stem_

Essentially, we wish to check whether our estimates of K_stem_ vary by less than a factor of 10 from literature values, whereby, for reference, a magnitude of 1 and 2 equate to actual values of 10 and 100 (10^1^ and 10^2^). In Shelly mango, log_10_K_stem_ is relatively similar between the 90th (1.03) and the 80th (0.92) percentile in the rainfall and irrigation period, respectively. This is consistent with the magnitude of K_s_ [kg s^−1^ MPa^−1^] in Cogshall (1.16), Irwin (1.10), and Kensington Pride (1.01) varieties, and slightly greater than José (0.61) (derived from Normand et al. (2008)). In Navel orange, log_10_K_stem_ has an order of magnitude of 0.13 in the 80th percentile rainfall period, which is close to an order of magnitude difference from K_s_ of 1.08 – 1.15 reported for well-watered Shamouti oranges grown in the same region as this study (Cohen et al., 1987). While *K*_*stem*_ could not be directly calculated as factor *a*_1_ is not known, we found that log_10_K_stem_ remains closely within an order of magnitude to literature values of *K*_*stem*_ for each species; this, in addition to the relatively close estimates in reductions in stem diameter to previously estimated MDS and ΔW in similar species and trees experiencing heatwaves, we conclude that the measurements for StWC and changes in StWS due to a heatwave seem reasonable.

## Supplementary material

### S1: Derivation of relationship between StWS and CHU

Transpiration *E* [mmol m^−2^ s^−1^], canopy conductance *g*_*c*_ [mmol m^−2^ s^−1^], VPD [kPa] and atmospheric pressure P [kPa] are commonly related by

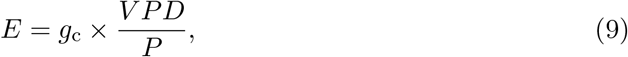

where *g*_*c*_ [mmol m^−2^ s^−1^] is the canopy conductance, and VPD is normalized by atmospheric pressure *P* [kPa], so that *E* and *g*_*c*_ have the same units.

where *g*_*c*_ can be be approximated to vary linearly with whole-plant leaf-specific hydraulic conductivity *K*_*L*_ [mmol m^−2^ s^−1^ MPa^−1^] (Hubbard et al., 2001) as

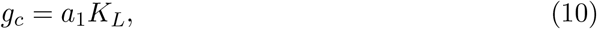

where the slope *a* is lower (higher) for isohydric (anisohydric) responses. ^1^ Substituting Eq. (10) into (9) yields

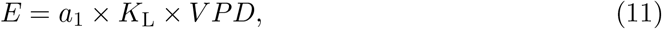

where atmospheric pressure *P* is taken to be constant, and thus is incorporated into the slope as *a*_1_. log(*a*_1_) has a magnitude of 1.5±0.5 (and *a*_1_ units are MPa Ψ_*stem*_ *atm*^−1^). ^2^.

We now convert the transpiration rate *E* into tree volume of transpired water *Ê* over a period *T*. We do this by integrating Eq. (11) over leaf area and over time, yielding

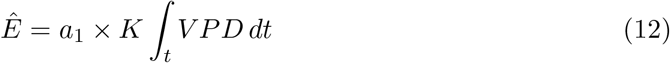

where the magnitude of *a*_1_ is unchanged due to the equal operations applied to both sides of the equation.

The total mass of transpired water in a day (*Ê*_*plant*_) must be equal to the total mass of water extracted from the storage pools available to the tree. This consists of water stored in the leaf (LWS), in the stem (StWS), and in the soil (SWS):

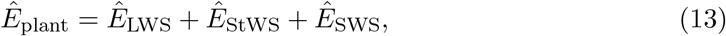

In the Mediterranean, it has been demonstrated that StWS acts as a buffer in the morning to midday hours when transpiration is high (Preisler et al., 2022). StWS diurnal flux drops during peak transpiration demand, and remains low until it is later restored by root uptake of SWS. We theorize, then, that in an extreme VPD event, this buffering property is exacerbated as it is stretched longer into the day, and that recharge from SWS, which becomes slower as transpiration reduces, will be insufficient to fully replenish StWS before it begins to act again as a buffer in the next day. In essence, there will be a net reduction in StWS within the period of an extreme VPD event.

Neglecting LWS as it is significantly less than the contributions from StWS and SWS, we quantify the relationship between water from storage pools used for transpiration with VPD by expanding equation XX with equation XX:

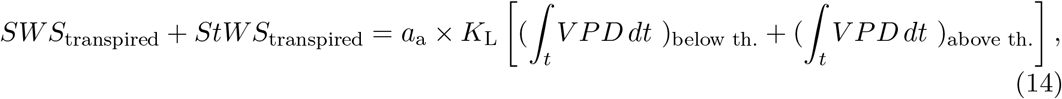

In a given day, if VPD remains below a high percentile threshold (i.e. a ‘regular’ day), then SWS eventually contributes the entire water budget since StWS from early-day buffering is replenished by SWS (Δ*StWS*_*day*_ = 0). Therefore, we first propose that a heatwave be identified by extreme VPD above the percentile threshold that leads to a net reduction in StWS within the day(s) of the event:

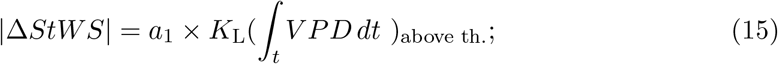

and that, second, the magnitude of the heatwave be quantified by the final term in the equation which incorporates both the intensity and duration of the extreme VPD event. We named this term the Cumulative Heatwave Unit (CHU), measured in kPa h:

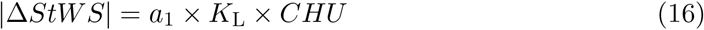

*Ê*_stem_ experiences resistance *K*_stem_

### S2: ag, the level of agreement in pairing extreme VPD events with negative StWC trend rate

While agreement generally ranges between 88-100% across percentiles and species, there are incidents (as explained in Section 2.6) in which there is a negative StWC trend rate with no corresponding VPD event, affecting ag. These may be explained as instances when StWC use is not related to transpiration (Preisler et al., 2022), or that transpiration demand was relatively—but not significantly—higher than a typical diurnal flux and was therefore carried forward during the de-trending step. The change in StWC trend in these instances was found to be negligible. Oppositely, the occurrence of statistically high VPD with no coincidental negative StWC trend rate was exceedingly rare, but still incorporated into ag.

### S3: log_10_K_stem_ (Table 4)

**Table 4:**
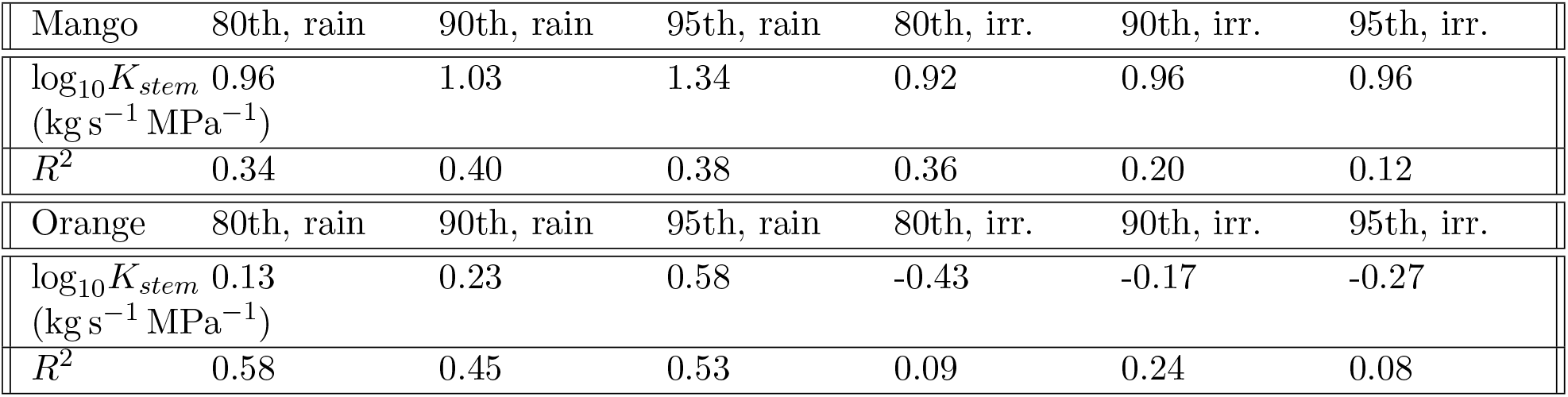
The magnitude of *K*_*stem*_ in mango relative to percentile threshold and period. The magnitude is calculated from the slope of the regression curves in figure 5 (log(slope/a1)).

### S4: Results of applied mixed linear model, grouped by tree label (Table 5)

The test was applied to the default case of species=Mango, season=Irrigated, percentile=80th. From this table, we see that the slope of the default case (0.0115) is statistically significant (p=0.000), and that the correction to the default slope by species=orange is negative (−0.0112) and statistically significant (p=0.000). This indicates that the slope of the orange response of StWS to CHU is lower than mango (slope=0.003), meaning that the orange trees from the study lose significantly less StWS as a response to heatwaves than the mango trees. There appears to be a significant effect by season on heatwave events (p=0.051), however the effect from VPD percentile is unclear (significant between the 80th and 95th percentile, p=0.008; insignificant between the 80th and 90th percentile, p=0.277). It is possible that were the study period to be extended to capture more events in the dry season, or to collect events from more than one year, that the statistical significance of season and VPD percentile may become stronger.

**Table 5:**
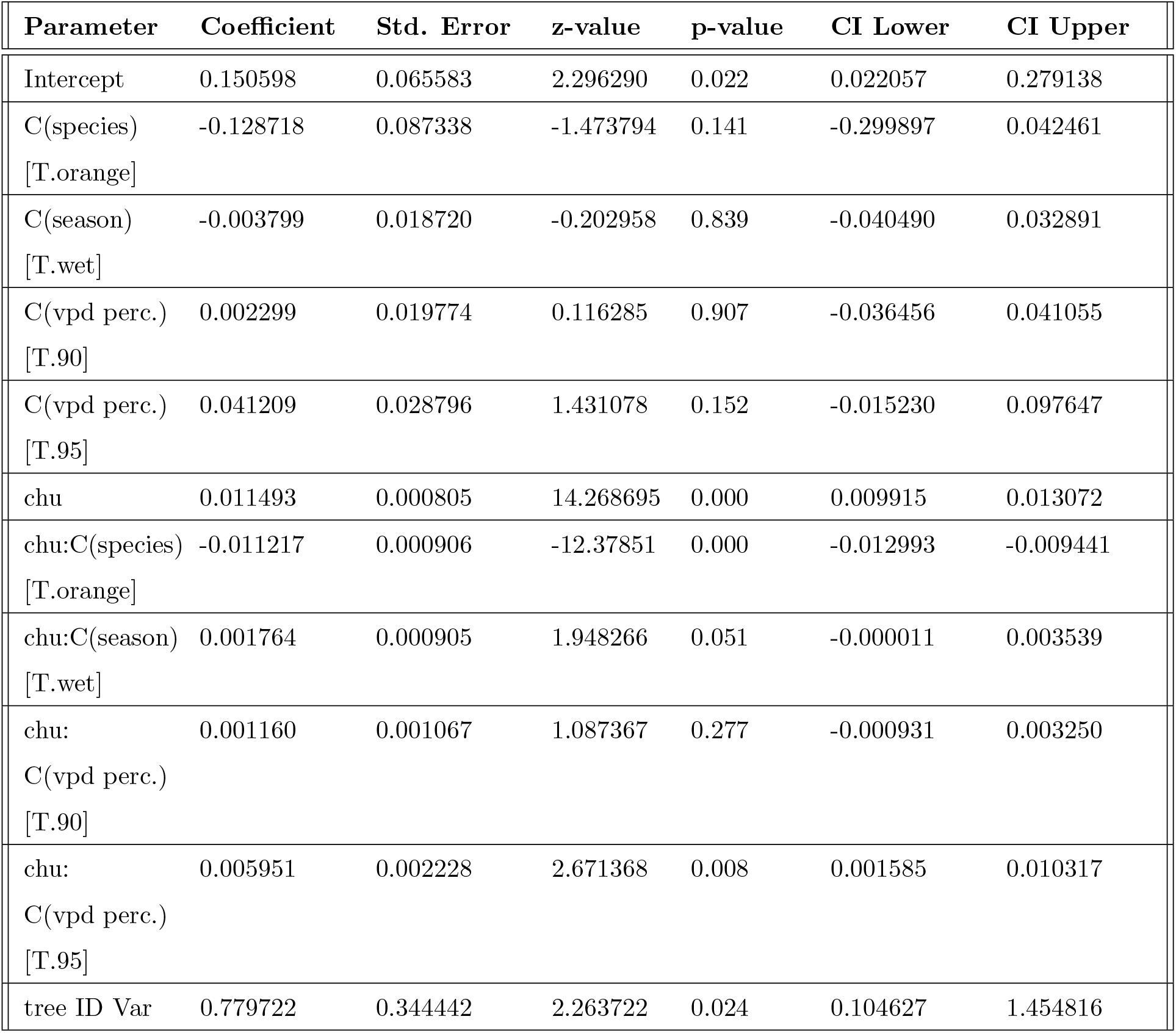
Mixed linear model: statistical significance of heatwave events by random effect of species, percentile threshold and season.

## Footnotes

1 *g*_*c*_ varies linearly with *K*_*L*_ for species with isohydric budgeting; the same is true for species with an anisohydric water budget, however *g*_*c,max*_ peaks before *K*_*L,max*_, creating a plateau towards *K*_*L,max*_.

2 In pine, *g*_*c*_ was found to be proportional to *K*_*L*_ by a factor of 33 (Hubbard et al., 2001), however this value can range higher for anisohydric species or for more isohydric species

## Notes

### Competing Interest Statement

The authors have declared no competing interest.

